## Supplemental Table 1 for "A novel RNA virus, *Macrobrachium rosenbergii* Golda virus (MrGV), linked to mass mortalities of the larval giant freshwater prawn in Bangladesh"

**Table S1**: Illumina MiSeq sequencing and assembly statistics.

| Library | Raw Reads | | rnaSPAdes Assembly | | | | | | IVA Assembly | | | | | Read Mapping | |
|---|---|---|---|---|---|---|---|---|---|---|---|---|---|---|---|
| | Paired reads (bp) | N50 | Number of contigs assembled | Minimum size (nt) | Maximum size (nt) | Average size (nt) | Contigs mapped to MrGV genome | Average Coverage (x) | Number of contigs assembled | Minimum size (nt) | Maximum size (nt) | Average size (nt) | Contigs mapped to MrGV genome | Paired reads mapped to MrGV genome | Average Coverage (x) |
| 1 | 3,453,843 | 133 | 4,713 | 139 | 11,163 | 643.3 | 11 | 0.2548 | 2 | 1,076 | 1,146 | 1,111 | 0 | 1,226 | 5.37 |
| 2 | 3,545,471 | 155 | 8,170 | 132 | 9,515 | 696 | 32 | 2.5333 | 20 | 869 | 3,393 | 1,727 | 0 | 49,594 | 269.28 |
| 3 | 5,867,622 | 209 | 19,533 | 131 | 14,156 | 674 | 50 | 5.04 | 14 | 937 | 14,858 | 4,457 | 2 | 248,277 | 1462.64 |
| 1+2+3 | 12,866,936 | 176 | 38,826 | 129 | 19,260 | 680 | 81 | 7.2555 | 13 | 735 | 29,110 | 5,046 | 1 | 299,141 | 1629.76 |
