## Supplemental Table 2 for "A novel RNA virus, *Macrobrachium rosenbergii* Golda virus (MrGV), linked to mass mortalities of the larval giant freshwater prawn in Bangladesh"

**Table S2:** Accession numbers and names of nidoviruses used for phylogenetic analyses.

| Taxonomic Classification | Species | Acronym | Accession number |
|---|---|---|---|
| *Arteriviridae* | Equine arteritis virus | EAV | X53459.3 |
| *Arteriviridae* | Kibale red-tailed guenon virus 1 | KRTGV | JX473849.1 |
| *Torovirinae* | Ball python nidovirus 1 | BPNV | KJ541759.1 |
| *Torovirinae* | Bovine torovirus | BRV | AY427798.1 |
| *Mesoniviridae* | Mesonivirus 2 | MenoV | JQ957873.1 |
| *Mesoniviridae* | Alphamesonivirus 1 | NDiV | DQ458789.2 |
| *Mesoniviridae* | Turrinivirus-1 | - | KX883629.1 |
| *Mononiviridae* | Planarian secretory cell nidovirus | PSCNV | MH933735.1 |
| *Coronavirinae* | Rousettus bat coronavirus HKU9 | Ro-BatCoV_HKU9 | EF065513.1 |
| *Coronavirinae* | Severe acute respiratory syndrome-related coronavirus | SARS-CoV | AY274119.3 |
| *Coronavirinae* | Severe acute respiratory syndrome coronavirus 2 | SARS-CoV-2 | NC_045512.2 |
| *Euroniviridae* | Charybnivirus-1 | - | KX883628.1 |
| *Euroniviridae* | Decronivirus-1 | - | KX883636.1 |
| *Euroniviridae* | Paguronivirus-1 | - | KX883627.1 |
| *Roniviridae* | Gill-associated virus | GAV | AF227196.2 |
| *Roniviridae* | Yellow head virus | YHV | EU487200.1 |
| *Astroviridae* | Chicken astrovirus | CAstV | MK746105.1 |
| *Astroviridae* | Canine astrovirus | CaAstV | NC_026814.1 |
